# Single-molecule Mechanostructural Fingerprinting of Nucleic Acid Conformations

**DOI:** 10.1101/2025.05.08.652980

**Authors:** Prakash Shrestha, William M. Shih, Wesley P. Wong

## Abstract

Understanding the three-dimensional structure and mechanical response of biomolecules is key to uncovering their molecular mechanisms, particularly in contexts where force plays a regulatory role. Structural methods such as X-ray crystallography, cryo-electron microscopy, and NMR spectroscopy provide high-resolution conformational data, while single-molecule force spectroscopy reveals mechanical properties—but these approaches are rarely integrated. A more comprehensive understanding of structure-function relationships, including non-equilibrium conformations and transitions under force, calls for methods capable of simultaneously resolving structural and mechanical properties at the single-molecule level. To meet this need, we present a DNA nanoswitch calipers platform capable of both measuring multiple intramolecular distances and mechanically unfolding individual biomolecules along defined axes. Using human telomeric DNA G-quadruplexes as a model system, we mapped distances between labeled sites to distinguish conformational states and performed directional unfolding to characterize mechanical stability along defined axes. This integrative approach revealed subtle conformational and mechanical differences, showcasing DNA nanoswitch calipers as a modular, broadly applicable approach for mechanostructural analysis of complex biomolecular systems.

## Introduction

Under physiological conditions, biomolecules experience mechanical stresses that can alter their conformations, drive structural transitions such as unfolding or bond dissociation, and in turn modulate their functions^1,2^. These responses often depend strongly on the direction of the applied force, highlighting the anisotropic nature of molecular mechanics^3–5^. Yet most experimental approaches isolate structure from mechanics, making it difficult to link force-induced conformational changes to function. Structural biology techniques such as cryo-EM, X-ray crystallography, and NMR spectroscopy provide high-resolution insights into molecular structures^6–8^, while single-molecule force spectroscopy methods like AFM and optical tweezers probe their mechanical properties and stability^9–12^. These approaches offer valuable, complementary perspectives, though each has its limitations. Structural techniques often fail to capture mechanical properties, molecular heterogeneity and dynamics^13^, while single-molecule force studies generally provide limited structural insight, with distances and mechanical stability typically only measured along a single axis^14,15^, and labor-intensive engineering of multiple constructs generally needed to assess multi-directional mechanical stability.

To address these gaps, an integrated approach is needed—one that combines structural and mechanical analyses of folded biomolecules. A technique capable of measuring multiple distances in a biomolecule to determine its 3D shape while enabling directional unfolding along various coordinates would provide a more comprehensive framework for investigating biomolecular mechanostructural properties, and their roles in biological processes. Here, we present an advanced DNA nanoswitch caliper (DNC) approach^16,17^ that enables determination of both 3D shape and mechanical stability of folded biomolecules at the single-molecule level. DNC is a mechanically reconfigurable DNA device, developed by our labs, that uses single-molecule force spectroscopy to map multiple distances by mechanically grabbing and releasing single-stranded DNA handles labeled at specific residues of a target biomolecule (Figure 1A). While earlier realizations of DNC measured structural distances, here we expand the platform to enable directional mechanical unfolding, integrating structural and mechanical characterization within a unified single-molecule framework.

**Figure 1.**
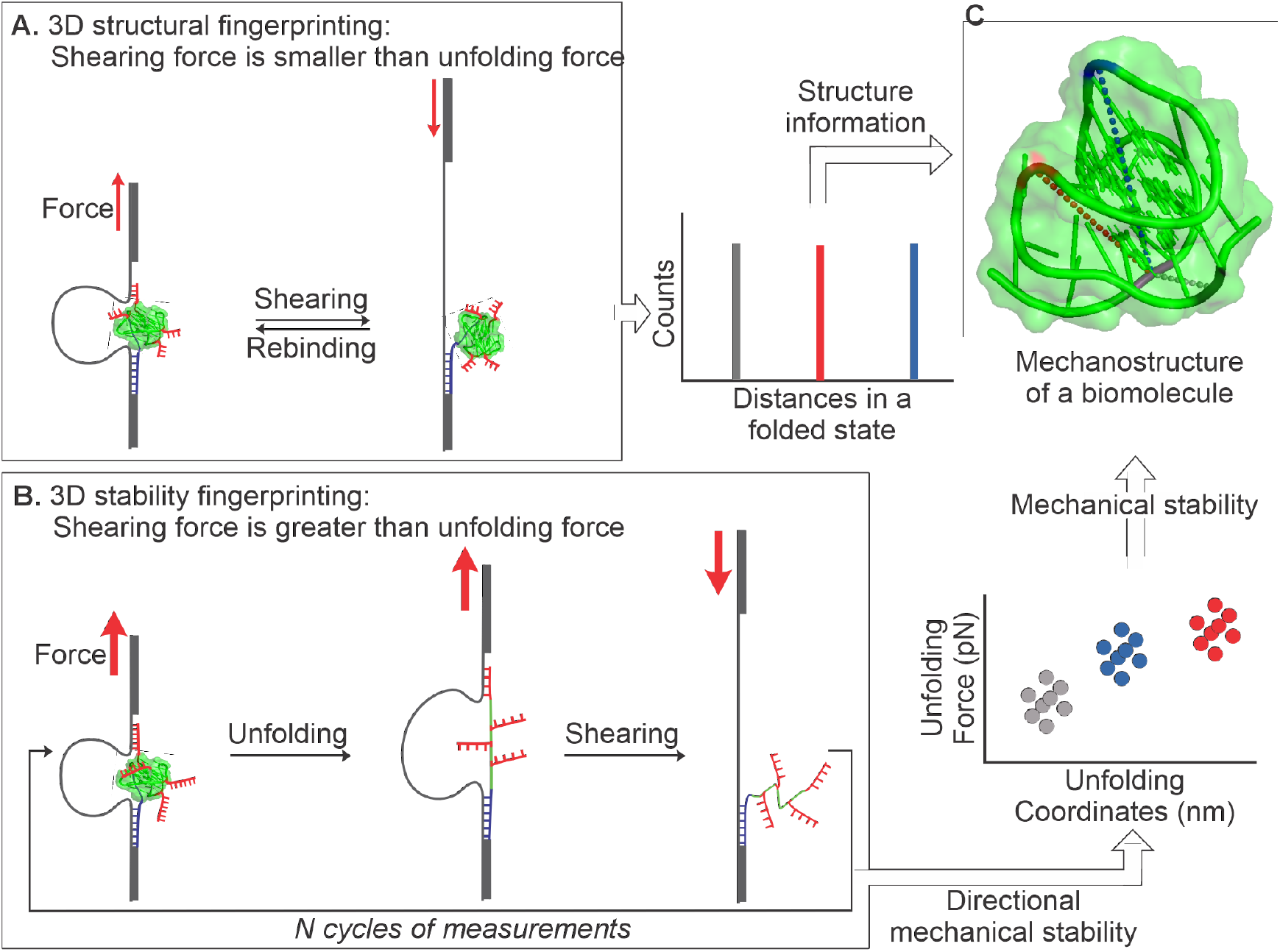
Schematic overview of mechanostructural fingerprinting using DNA nanoswitch calipers (DNC). **(A)** Design of DNC for measuring multiple distances within a folded biomolecule. Shorter shearing handles are used to unloop the DNC before the structure unfolds. **(B)** Design of DNC for measuring the mechanical stability of a folded biomolecule along different pulling coordinates. Stronger shearing handles allow the structure to be unfolded prior to DNC unlooping, enabling directional unfolding along multiple axes. **(C)** Structural information from multiple distances in the folded state (A) combined with directional mechanical stability data (B) enables construction of a mechanostructural fingerprint of the folded biomolecule. Dotted lines depict the measured distances, and color coding reflects mechanical stability along different axes.

As a proof of concept, we applied our approach to the human telomeric DNA G-quadruplex, a biologically significant yet labile structure formed by Hoogsteen H-bonding between guanine bases and stabilized by specific metal cations. These structures can adopt distinct conformations, depending on nucleotide sequence, buffer conditions and the identity of the stabilizing cations^18,19^. Understanding the mechanical stability of these conformations is biologically and physiologically important, as G-quadruplexes can act as mechanical roadblocks that must be unfolded by motor proteins to allow continued replication or transcription. Similarly, RNA G-quadruplexes are known to regulate translation, and telomeric G-quadruplexes can regulate telomerase activity and influence telomere length^12,20–22^. We demonstrate that DNC enables the determination of both G-quadruplex conformation, through measurement of multiple intramolecular distances in the folded state (Figure 1A), and anisotropic mechanical stability, through directional unfolding along multiple axes (Figure 1B), producing a distinct mechanostructural fingerprint (Figure 1C). This study establishes DNC as a simple yet modular platform for single-molecule mechanostructural analysis of biomolecules and their complexes.

## Materials and methods

### Materials

The pET-26b (+) plasmid was received from EMD Millipore Sigma. All the DNA oligonucleotides (see sequences in Supplementary Table 1) were purchased from Integrated DNA Technologies (IDT). The enzymes required to synthesize DNC constructs were purchased from New England Biolabs. Chemicals such as 1-Ethyl-3-(3-dimethyl aminopropyl) carbodiimide (EDC), and N-hydroxysuccinimide (NHS) ester were purchased from Sigma Aldrich. The carboxylated silica beads (3 µm) were purchased from Microspheres-Nanospheres.

### Site-specific labeling of G-quadruplex with single-stranded DNA handles

The single-stranded DNA oligo with a human telomeric sequence (TTAGGG)_4_ that had an azide functional group in the first thymine base of each loop sequence (TTA) was incubated with the DNA handle labeled with DBCO at the 5′ end. The reaction mixture was prepared in 1x PBS buffer and sealed with paraffin film and incubated for more than 18 hours in no direct light. The completion of the reaction was tested in PAGE electrophoresis assay and the fully labeled target was purified from the gel (Supplementary Figure 1).

### Synthesis of DNA Nanoswitch Caliper (DNC) constructs

Two different DNA nanoswitch calipers constructs were synthesized, one incorporating a short loop and the other a long loop, following previously described methods^16,17^. Briefly, the long dsDNA handle (2820 bp) used in both constructs was prepared by polymerase chain reaction (PCR) amplification of a specific region of the pET-26b (+) plasmid. The forward primer contained a BsaI restriction site, and the reverse primer was labeled with dual biotins at the 5′ end (see sequences in Supplementary Table 1). The PCR product was purified using a PCR purification kit (Qiagen) and then digested with BsaI restriction enzyme to create sticky ends for ligation with the other fragment of the DNC construct. The other DNC fragment was prepared by annealing the loop oligonucleotide (short or long loop), 5′-digoxigenin modified oligonucleotide, and other required strands (Supplementary Table 1) at equimolar concentration with a splint oligonucleotide added at 10-fold molar excess. The final DNC construct was synthesized by ligating this annealed fragment to the dsDNA handle using T4 DNA ligase (New England Biolabs) at 16 °C for 16 h, followed by heat deactivation of the ligase at 65 °C for 20 min. The fully ligated product was purified using agarose gel electrophoresis.

### Single-molecule distance and unfolding force measurements using optical tweezers

We used home-built dual-trap optical tweezers for all DNC measurements^23^. The short-loop DNC was used to measure distances in the folded G-quadruplex, while the long-loop DNC was used to unfold the structure along multiple different pulling axes. All DNC measurements were performed in tris buffered saline (TBS, 150 mM NaCl or 150 mM KCl, 10 mM tris buffer, pH 7.4) with 0.1

% (w/v) Roche Blocking solution (Roche) at room temperature (23 °C). First, 5 µL of DNC construct (∼100 pM) was incubated with 0.5 µL of a 1% (w/v) solution of streptavidin-coated silica beads (3.0 µm diameter) for 30 min to immobilize the DNC construct on the surface of the beads through binding of biotin and streptavidin. The DNC-functionalized beads were washed twice by centrifugation to remove unbound constructs. For an end-to-end distance measurement in a folded G-quadruplex, the target DNA consisting of a human telomere G-quadruplex sequence sandwiched between a grabbing handle sequence at the 5′ end and a shearing handle sequence at the 3′ end was incubated with the DNC immobilized beads for 30 minutes. Then, the beads were washed to remove excess target DNA oligonucleotides. To measure multiple distances in an individual G-quadruplex, G-quadruplex labeled with shearing handles in each loop site (as prepared above), was incubated with the DNC immobilized beads for 30 minutes, followed by washing to remove the unbound targets.

The sample cell was prepared as described previously^16,17^. Briefly, a double-sided Kapton tape (1 mm, DuPont) was sandwiched between a glass slide and a cover glass (prewashed with 70% ethanol and dried with argon flow). Two side channels were created by cutting the Kapton tape before sandwiching for injecting DNC immobilized beads and anti-digoxigenin-coated beads. The channel was passivated with Roche Blocking solution, (1 % w/v) for ∼30 min and then flushed with the experimental buffer before injecting the beads. The DNC immobilized beads (1 µL) were injected through one side-channel and anti-digoxigenin coated silica beads (1 µL of the 0.1% (w/v)) were injected through the next side-channel. The channel openings were then sealed with grease to prevent dry-up of the sample during measurements. In dual-trap optical tweezers, the two different beads were trapped in separate laser foci and a tether was formed by binding digoxigenin at the end of the immobilized DNC with an antidigoxigenin-coated bead as shown in Figures 2-5. The tethered DNC was confirmed as a single molecule by estimating the contour length of the whole caliper (∼1 µm) and by a single breakage of the tether or the observation of a ∼65 pN plateau in force due to mechanical overstretching of dsDNA. The distance measurement on the folded G-quadruplex was carried out by shearing the handle at a low force (∼10 pN, which is less than the approximate unfolding force of the G-quadruplex of ∼21 pN) and then jumping to a force of ∼0 pN to let the shearing handle rebind. By repeatedly cycling between force levels, extensive data was collected from each individual caliper. For directional unfolding of the G-quadruplex, a G-quadruplex labeled with the longer shearing handles (see Supplementary Table 1) was used. Slow ramping of the force (∼6 pN/s) up to ∼45 pN was used to probe the unfolding of the G-quadruplex, followed by a rapid jump to a higher force to unloop the DNC to enable identification of the attachment site on the construct. Data was recorded at 1400 Hz using a custom LabVIEW 2013 software (National Instruments Corporation, Austin, TX).

**Figure 2.**
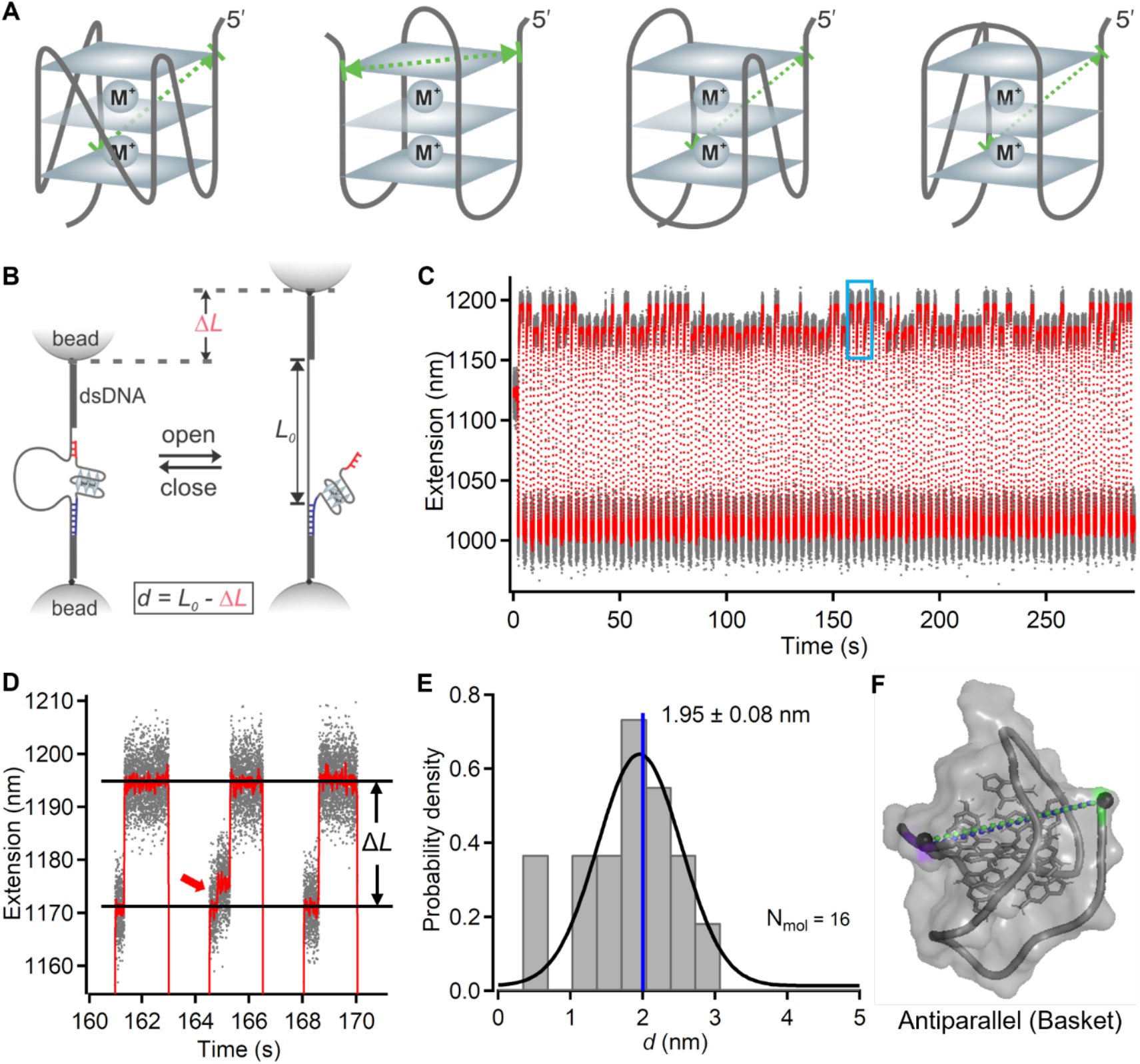
Single-molecule measurements of 5′-3′ distance in a folded G-quadruplex in sodium buffer. **(A)** Schematic structures of four representative G-quadruplex conformations. Left to right: Parallel, Antiparallel, Hybrid 1, and Hybrid 2. Green dotted lines with arrowheads indicate the end-to-end distance for each structure. **(B)** Schematic depiction of the measurement strategy using DNA nanoswitch calipers and optical tweezers. A longer handle at the 5′ end (purple) serves as a strong grabbing handle, while a shorter handle at the 3′ end (red) serves as a weak shearing handle. Unlooping of the DNC due to the shearing of the shorter handle is recognized as a change-in-length (Δ*L*). The absolute distance (*d*) is calculated by subtracting Δ*L* from the loop length of the DNC (*L*_*0*_). **(C)** Representative extension vs. time graph showing Δ*L* measurements obtained by jumping the force between ∼10 pN and ∼0 pN in repeated cycles. Gray trace shows raw data, while the red trace indicates sliding-window averaged data (window size = 200). **(D)** Zoomed-in view of the highlighted section in (C), indicated by the blue rectangle, showing repeated distance measurements and occasional unfolding events (red arrow) during force cycles. **(E)** Histogram of per-molecule averaged 5′-3′ (end-to-end) distances for 16 G-quadruplexes. Plus-minus values indicate the standard error of the mean. The blue vertical line indicates the NMR-determined distance for the antiparallel (basket) conformation. N_mol_ denotes the number of measured molecules. **(F)** Comparison of the 5′-3′ distance measured by DNC (green dotted line) and that reported by NMR (PDB 143D; blue dotted line) for the antiparallel (basket) conformation showing strong agreement.

### Data analysis

For all single-molecule measurements, data was acquired using custom LabVIEW software and analyzed with custom MATLAB scripts. Statistical analyses and graphing were performed using IGOR Pro. G-quadruplex unfolding was identified by a sudden change in extension during slow ramping of the force (Figure 5), whereas DNC unlooping was determined by a discrete change-in-length (*ΔL*) during the holding period at the shearing force (Figure 2-5). To convert the observed *ΔL* from DNC unlooping into absolute distance *d*, we subtracted *ΔL* from the effective loop length *L*_*0*_, i.e., *d* = *L*_*0*_ - *ΔL. L*_*0*_ at the measurement force was determined by measuring *ΔL* for analytes of known length and performing a linear regression to extrapolate the change-in-length for a zero-length analyte, i.e., the offset of the linear fit is *L*_*0*_. However, for multiple distance and directional unfolding measurements, we interpreted results directly in terms of *ΔL, as L*_*0*_ varied across configurations due to the presence of additional linkers in the loop-region shearing handles as compared to those at the 3′-end.

### Circular Dichroism Spectroscopy

Human telomere DNA oligonucleotide (Supplementary Table 1) samples were prepared at 5 µM in 10 mM Tris buffer (pH 7.4) containing either 150 mM NaCl or 150 mM KCl. Samples were first heated at 95°C for 5 min, then rapidly cooled in an ice bath, followed by incubation at room temperature for up to an hour. CD spectroscopy was performed in a 1 mm quartz cuvette at room temperature with a JASCO J-1500 spectropolarimeter. Scans were performed from 200 nm to 320 nm at a rate of 100 nm/min. Reported spectra represent the average of the recorded data using a window size of 100. The spectrum of the corresponding buffer was used for baseline correction.

## Results and Discussion

### Measurement of the distance between 5′ to 3′ ends in a folded G-quadruplex structure in different ionic conditions

DNA G-quadruplexes are polymorphic, with conformations governed by the type of stabilizing cations and the nucleotide sequences^24–26^. There are different types of DNA G-quadruplex conformations such as parallel, anti-parallel, and mixed-type hybrid-1 and hybrid-2. Depending on the folded G-quadruplex conformation, the 5′ to 3′ end-to-end distances can vary (Figure 2A).

We confirmed that the target human telomere sequence adopts distinct conformations in sodium and potassium buffers using circular dichroism spectroscopy (Supplementary Figure 2). To measure the end-to-end distance in the folded state at the single-molecule level, we designed a short (9 base) DNA handle at the 3′ end that shears open the DNA nanoswitch caliper at ∼10 pN, below the ∼20 pN unfolding force of the G-quadruplex^11,17^. First, we measured the end-to-end distance of the folded G-quadruplex in sodium buffer (10 mM Tris pH 7.4, 150 mM NaCl) as shown in Figure 2B. In repeated force cycles, we jumped the force between two different levels— ∼10 pN to shear open the DNC, indicated by a sudden change in extension (Δ*L*) (Figure 2 B&C), and ∼0 pN to allow the handle to rebind, enabling repeated distance measurements within the same molecule. At the shearing force, G-quadruplex unfolding sometimes occurred prior to DNC unlooping, as indicated by the red arrow in Figure 2D, and hence the histogram of the change-in-distance measured from several molecules showed three major populations: ∼6.0 nm due to unfolding of the G-quadruplex, ∼20.0 nm due to unlooping from the unfolded G-quadruplex, and ∼26.0 nm due to unlooping from the folded G-quadruplex (Supplementary Figure 3). By subtracting the change-in-distance Δ*L* associated with DNC unlooping in the folded G-quadruplex state from the loop length, we obtained the distribution of the measured end-to-end distance of the folded G-quadruplex, which indicated a single population centered at ∼2.0 nm (Figure 2E). This distance of ∼2.0 nm measured using DNC matches that of the antiparallel conformation reported by NMR^27,28^, confirming that the folded G-quadruplex adopts this conformation in these sodium buffer conditions (Figure 2F). Single-molecule FRET measurements have also reported the antiparallel conformation as a major population in sodium buffers but also observed a certain extent of conformational heterogeneity suggesting that multiple interconvertible states could exist under different conditions^29^.

On the other hand, when we switched from sodium to potassium buffer (10 mM Tris pH 7.4, 150 mM KCl), the measured 5′-3′ (end-to-end) distance distribution shifted to ∼1.0 nm, indicating that the G-quadruplex adopted one of several alternative conformations (Figure 3). Three candidate structures, each with similar end-to-end distances, have been reported in NMR studies^30,31^ (Supplementary Table 2). Distinguishing between these conformations requires the measurement of multiple distances within the folded G-quadruplex.

**Figure 3.**
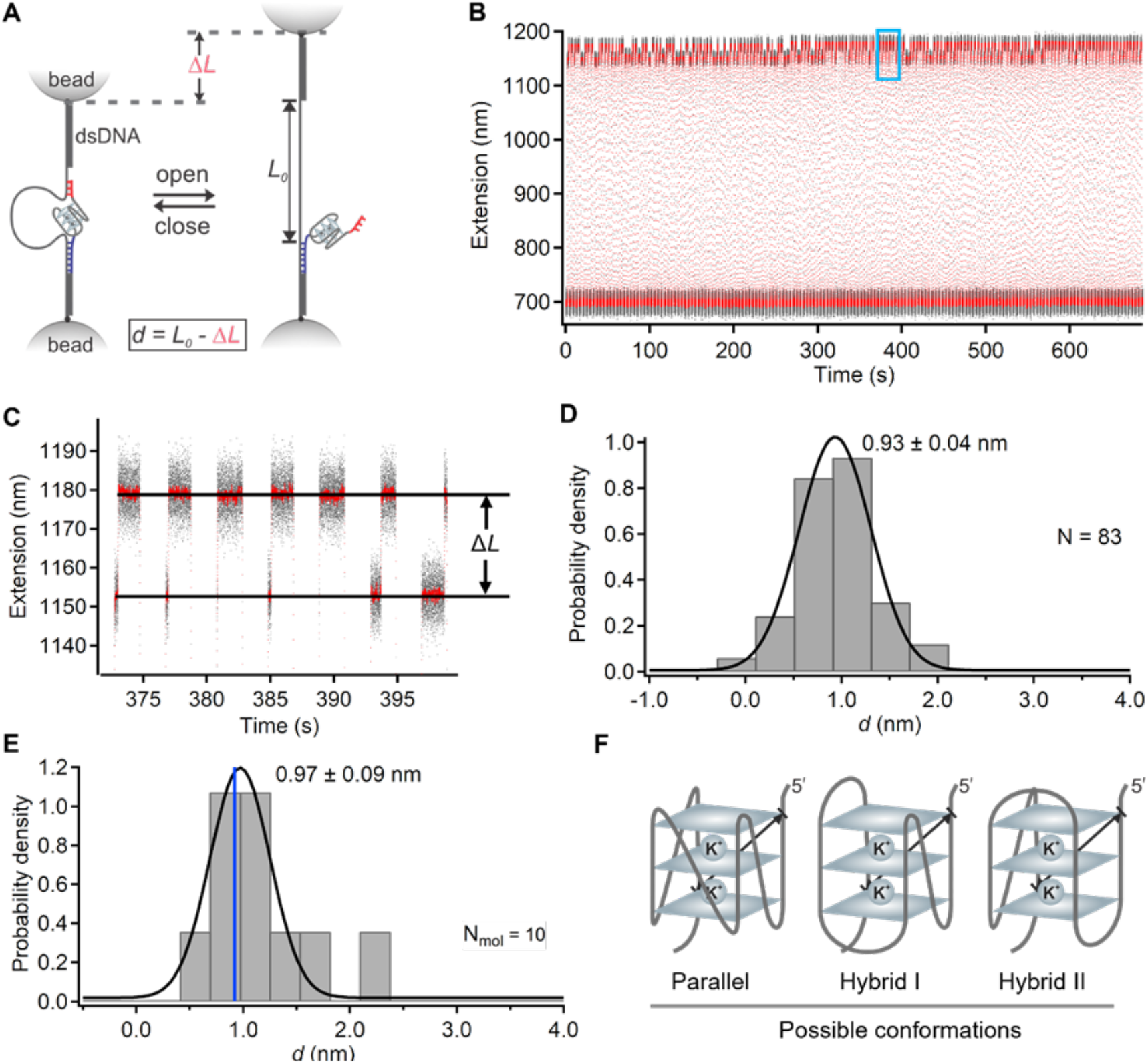
Single-molecule measurement of 5′-3′ distance in a folded G-quadruplex in potassium buffer. **(A)** Schematic depiction of the same measurement strategy shown in Figure 2B, now applied in potassium buffer. A longer grabbing handle at the 5′ end (purple) and a shorter shearing handle at the 3′ end (red) are used to measure 5′-3′ distance in a folded G-quadruplex via DNC unlooping. **(B)** Representative extension vs. time graph showing Δ*L* measurements obtained by cycling the force between ∼10 pN to shear the red handle and ∼0 pN to enable handle rebinding. Gray trace shows raw data, while the red trace indicates sliding-window averaged data (window size = 200). **(C)** Zoomed-in view of the section indicated by the blue rectangle in (B) showing repeated distance measurements in a folded G-quadruplex. **(D)** Histogram of 5′-3′ (end-to-end) distances measured repeatedly in a single G-quadruplex. **(E)** Histogram of per-molecule averaged distances for 10 G-quadruplexes. Plus-minus values indicate the standard error of the mean from Gaussian fitting. The blue vertical line indicates the 5′-3′ distance from the NMR-determined structure. **(F)** The measured 5′-3′ distance is consistent with three possible conformations reported in structural studies.

### Measurement of multiple distances in a folded G-quadruplex in potassium buffer

Next, to resolve between the three potential conformations of the G-quadruplex in potassium buffer, we utilized the capability of DNA nanoswitch calipers to measure multiple distances within the same biomolecule^16,17^. A G-quadruplex with multiple shearing handles was prepared by labeling the first thymine base in each loop region using DBCO-azide click chemistry (Supplementary Figure 1). We measured pairwise distances between the 5′ end and each of the three loop-labeled handles, as well as the 3′ end of each folded structure. As before, we performed these measurements by cycling between a shearing force of ∼10 pN and a rebinding force of ∼0 pN to enable sampling of all four distances (Figure 4 A-E). The distribution of the change-in-distance (Δ*L)* values showed multiple peaks, as expected (Figure 4E, and Supplementary Figure 5). After per-molecule averaging, we found four distinct populations, indicating successful measurement of all four distances in the folded G-quadruplex (Figure 4F). To identify the most likely conformation, we compared the measured distances to those expected from published NMR structures and calculated the root-mean-squared deviation (RMSD) for each (Supplementary Table 2). Based on the lowest RMSD, the conformation of the G-quadruplex in potassium buffer is most consistent with the hybrid I (PDB 2HY9) conformation (Figure 2G). This assignment also matches previous reports^32^. These results demonstrate the capability of DNA nanoswitch calipers to mechanically probe and distinguish between conformations of folded biomolecular structures. We also note that both static and dynamic heterogeneity could be present within this population of molecules. In other words, individual molecules could adopt distinct conformations, or a single quadruplex could potentially interchange between different conformations over successive measurements. While this data set alone cannot definitively resolve these possibilities, in principle, single-molecule approaches should be well-suited to characterize such heterogeneity^33^.

**Figure 4.**
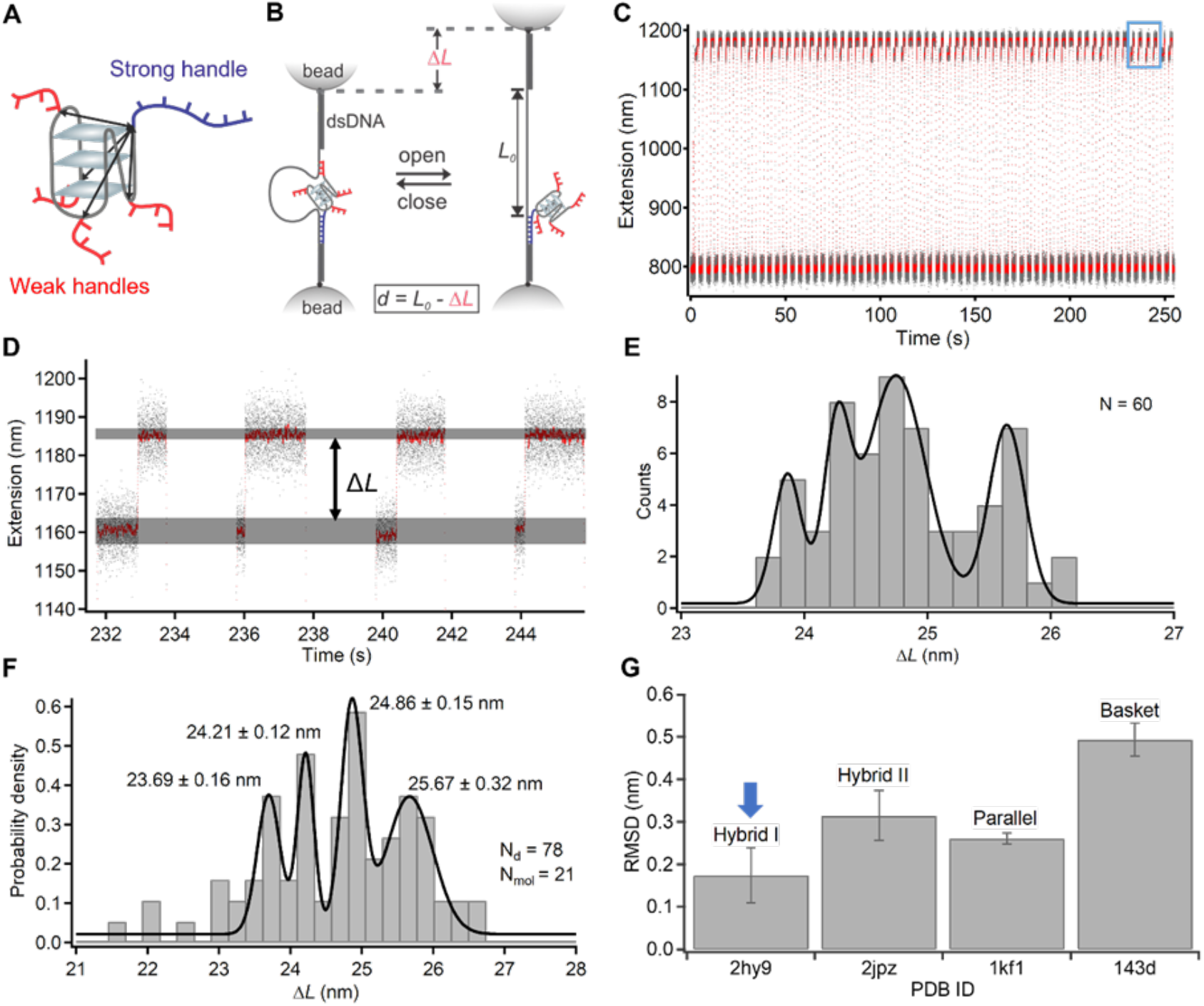
Measurement of multiple distances in a folded G-quadruplex in potassium buffer. **(A)** Schematic representation of the G-quadruplex structure showing the four potential pairwise distance measurements. One strong grabbing handle (purple) is positioned at the 5′ end, and four weak shearing handles (red) are attached at three internal loop positions and at the 3′ end. **(B)** Schematic of the DNC measurement strategy using optical tweezers. **(C)** Raw extension vs. time data showing the change in extension over multiple measurement cycles (gray); red trace indicates sliding-window averaged data (window size = 200). **(D)** Zoomed-in view of the region indicated in (C) (blue rectangle) showing unlooping events for different measured distances within the folded G-quadruplex. Horizontal gray bands indicate the states before and after DNC unlooping; the broader band in the unlooped state reflects variation due to rebinding to different shearing handles. **(E)** Histogram of change-in-distance (Δ*L*) values measured in a single G-quadruplex. **(F)** Histogram of per-molecule averaged Δ*L* values measured in 21 G-quadruplexes. Plus-minus values indicate sigma fitting parameters from the multiple Gaussian fit. **(G)** Root-mean-squared deviation (RMSD) analysis comparing measured Δ*L* values to those expected for four representative G-quadruplex conformations.

### Directional unfolding of individual G-quadruplex structures in sodium buffer

Mechanical unfolding using single-molecule techniques enables directional probing of molecular stability, distinct from the more global unfolding induced by chemical denaturants or changes in temperature. This approach has provided valuable insights into kinetic intermediates and folding pathways in biomolecules^34–37^. However, such measurements are often challenging, as single-molecule mechanical unfolding is typically performed along a single pulling direction per construct, requiring multiple different designs to interrogate direction-dependent behavior^14,35,38,39^. Here, we demonstrate how DNA nanoswitch calipers can be used to unfold individual G-quadruplex structures in multiple different directions. To enable this, we designed G-quadruplex constructs with longer shearing handles (16 bases) at the 3′ end (Supplementary Figure 6 and Supplementary Table 1), as well as in each loop region, labeling the first thymine base in each loop region using DBCO-azide chemistry. These longer handles were designed so that unlooping of the DNC would occur at a higher force than G-quadruplex unfolding, allowing us to observe first unfolding, then opening of the DNC to identify which handle was grabbed and know the direction of unfolding. During each experimental cycle, unfolding of the G-quadruplex was observed as a small, sudden change in extension (Δ*L*_unfold_) of ∼2-9 nm during a period of slow force ramping up to ∼45 pN. The force was then rapidly increased to >55 pN to shear open the DNC, producing a larger change in extension (Δ*L*) of ∼58-68 nm, which revealed the handle through which the G-quadruplex had been unfolded (Figure 5 A-D). We observed unfolding along three distinct pulling directions (5′-end to mid-loop, 5′-end to the 3′-end loop, and 5′-end to the 3′-end) as confirmed by the expected Δ*L*_unfold_ and Δ*L* values within each cycle (Figure 5E and Supplementary Table 4). As expected, we did not observe unfolding through the 5′-end loop, as the expected change in distance due to unfolding was below our detection threshold in this pulling direction (Supplementary Table 4). When comparing unfolding forces across different pulling directions, we found that pulling through the loops required a slightly higher force than pulling from the 5′-end to the 3′-end (Figure 5F). This may reflect differences in the alignment of Hoogsteen H-bonding relative to the pulling direction: unfolding from 5′ to 3′ proceeds along the stacking axis in an unzipping-like mode, whereas pulling through the loops engages the G-quadruplex at more oblique angles, which may require higher forces for disruption.

**Figure 5.**
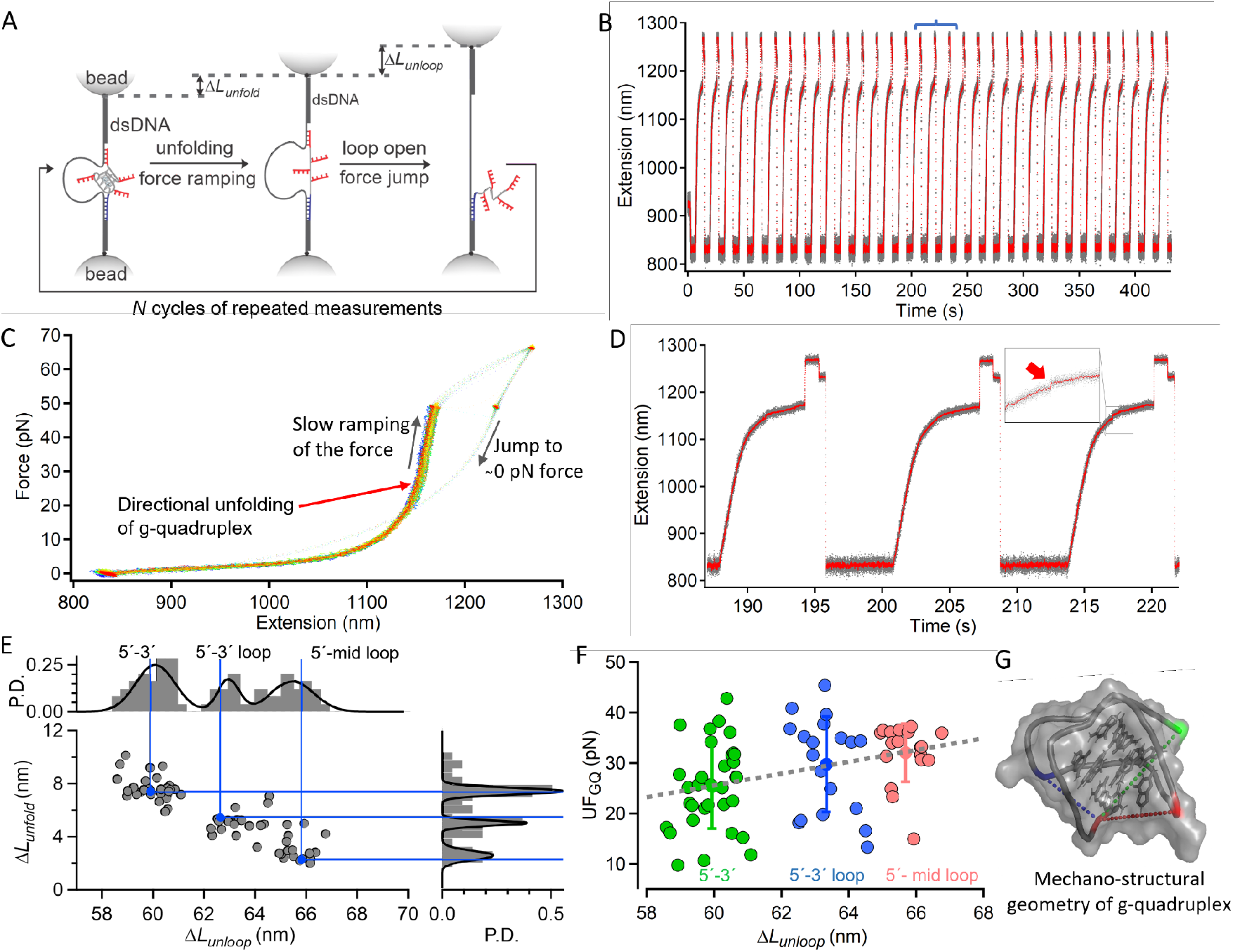
Directional unfolding of individual G-quadruplex structures along multiple pulling directions. **A)** Schematic representation of the design and experimental strategy to unfold the individual G-quadruplex through multiple directions. First step involves the slow ramping of force to probe the unfolding of the G-quadruplex, and the second step involves the jumping of force to shear-open the DNC to probe the site through which the G-quadruplex was unfolded. **B)** An experimental graph of change-in-extension over time. Gray trace indicates raw data, while red trace indicates sliding-window averaged data (window size=100). **C)** Overlapped force vs. extension curves from all the repeated measurements of the same G-quadruplex structure. A spread region in the plot is due to the unfolding of the G-quadruplex in several cycles. **D)** Zoomed in on a section indicated by a blue line in the extension vs. time plot in B) to show the G-quadruplex unfolding features (red arrow at the inset). **E)** Scatter plot of the change-in-extensions due to the unfolding of G-quadruplex and unlooping of DNC indicate the specific site via which the G-quadruplex was unfolded. Histogram of the change-in-distance due to unlooping of DNC (Δ*L*_unloop_) (top) and histogram of the change-in-distance due to unfolding of G-quadruplex (Δ*L*_unfold_) (right). Black solid curves depict the multiple peak Gaussian fitting. Blue lines indicate the expected change-in-distance due to unlooping (vertical) and unfolding of the G-quadruplex (horizontal). **F)** Correlation of the unfolding force of the G-quadruplex with the direction from which it was unfolded. Color codes are used to indicate the clustering of the unfolding force of G-quadruplex when pulling through different directions determined by measuring the change-in-distance due to unlooping of DNC. Gray dotted lines depict the linear fitting of the mean values of the corresponding clusters. **G)** Depiction of the mechanical stability of G-quadruplex along different coordinates on the G-quadruplex conformation in sodium ions reported by NMR (PDB 143D) and also matched with DNC measurement (Figure 2). Color-coded residues indicate the respective pulling directions as mentioned in (F) and dashed lines indicate the distance between those residues in the reported structure.

By combining the measurement of distances in the folded state (Figure 2) with directional unfolding data (Figure 5), we generated a mechanostructural fingerprint of the G-quadruplex conformation (Figure 5G), providing an integrated picture of its structure and mechanical anisotropy. Recent work by Yang et al.^5^ and Sedlak et al.^4^, demonstrate that mechanical stability in receptor-ligand complexes can strongly depend on tethering geometry, establishing that the direction of force application alone can modulate unbinding behavior. Our results extend this concept to folded nucleic acid structures, revealing that the direction of applied force can significantly influence unfolding behavior. Such directional sensitivity could impact how effectively G-quadruplex structures act as mechanical roadblocks during replication and transcription, and how telomerase engages telomeric DNA substrates. By enabling directional unfolding and multi-point distance measurements within the same molecule, DNA nanoswitch calipers overcome key limitations of previous single molecule approaches. This method will provide a powerful tool for probing mechanostructural relationships in nucleic acids and their protein complexes, enabling direct investigations into how structural conformations respond to biologically relevant mechanical forces.

## Author contributions

P.S., W.W., and W.S. conceived the project. P.S. and W.W. designed the experiments. P.S. conducted experiments and performed data analysis. All authors discussed the results and analysis, and contributed to the manuscript, with the initial draft written by P.S. and W.W.

## Supplementary data

Supplementary information is available online for this paper.

## Data availability

The data underlying this article will be made available by the corresponding authors upon reasonable request.

