## Supplementary Information for "Single-molecule Mechanostructural Fingerprinting of Nucleic Acid Conformations"

**Supplementary Table 1.** List of DNA oligonucleotides that are used to synthesize DNA Nanoswitch Calipers (DNC), target DNA oligonucleotides and DNA handles used to label the G-quadruplex. Purified DNA oligonucleotides were purchased from IDT.

| **Oligonucleotides** | **Sequences** |
|---|---|
| BsaI-forward primer | AAA AAA GGT CTC GGC ATG ATA GCG CCC GGA AGA GAG |
| Biotin-reverse primer | /52-Bio/GGG TTC GTG CAC ACA GCC CAG CTT |
| Loop_short | ATG CCG CGT AAC CGG TCC GAG ACT CTC ACT GTG AAC TGA GTT TTT TTT TTT TTT TTT TTT TTT TTT TTT TTT TTT TTT TTT TTT TTT TTT TTT TTT TTT TTT TTT TTT TTT TTT TTT TTT TTT TCA GCG TCC GCG TGT CAG TGT |
| Loop_long | ATG CCG CGT AAC CGG TCC GAG ACT CTC ACT GTG AAC TGA GTT TTT TTT TTT TTT TTT TTT TTT TTT TTT TTT TTT TTT TTT TTT TTT TTT TTT TTT TTT TTT TTT TTT TTT TTT TTT TTT TTT TTT TTT TTT TTT TTT TTT TTT TTT TTT TTT TTT TTT TTT TTT TTT TTT TTT TTT TTT TTT TTT CAG CGT CCG CGT GTC AGT GT |
| DNC1 | /5DigN/CGT CCC CAA ATC CGT ACC GGA TGC ACT GGC ACT GTG GGT GTG CGA CTT AAT TCC ATC CTG TTC CTG AGA CAA TAC AGC ACG ACC GCT TTC |
| DNC2 | /5Phos/TCT AAC TTA CAG AGC ATG GCG AAA GCG GTC GTG CTG TAT TGT CTC AGG AAC AGG ATG GAA TTA AGT CGC ACA CCC ACA GTG CCA GTG CAT CCG GTA CGG ATT TGG GGA CG |
| DNC3 | TCT CGG ACC GGT TAC GCG |
| Splint | AAG TTA GAA CAC TGA C |
| DNC4 | GAT GGC CCG CTG AAA GGG CAG TGT TTC CCA GCG CCC TTC CTG GTA TGC GGA TTC TTT CGG GAG ATA GTA ATT AGC ACC TCC ATC GTC AGA CGG CGT CTG TGG CTC CGG CCT GAA CAG TGA |

| | |
|---|---|
| DNC5 | /5Phos//TCT AAC TTA CAG AGC ATG GCT CAC TGT TCA GGC CGG AGC CAC AGA CGC CGT CTG ACG ATG GAG GTG CTA ATT ACT ATC TCC CGA AAG AAT CCG CAT ACC AGG AAG GGC GCT GGG AAA CAC TGC CCT TTC AGC GGG CCA TCG TGA TTT GAC TGT CTC CGG CCT TTC TCA CCC TTT TGA ATC TTT ACC TAC ACA TTA CTC AG |
| GH-GQ-LM | GCC ATG CTC TGT AAG TTA GAA CAC TGA CAC GCG GAC GCT GTT AGG G/iAzideN/T AGG G/iAzideN/T AGG G/iAzideN/T AGG GTT ACT CAG TTC A |
| GH-GQ-9bSH | GCC ATG CTC TGT AAG TTA GAA CAC TGA CAC GCG GAC GCT GTT AGG GTT AGG GTT AGG GTT AGG GTT ACT CAG TTC A |
| GQ-9bSH | /5DBCON/CTC AGT TCA |
| GH-GQ-9bSH-ext | /5Phos/CAG TGA G |
| Ext-splint | CAT GTC GAC TCA CTG TGA ACT GAG T |
| Ext-splint-removal | ACT CAG TTC ACA GTG AGT CGA CAT G |
| GQ-16bSH | /5DBCON/CTC AGT TCA CAG TGA G |

Sequences from left to right are in the 5´ to 3´ directions. Underlined sequence represents the human telomere sequence (TTAGGG)$_4$ that can fold into a G-quadruplex structure.

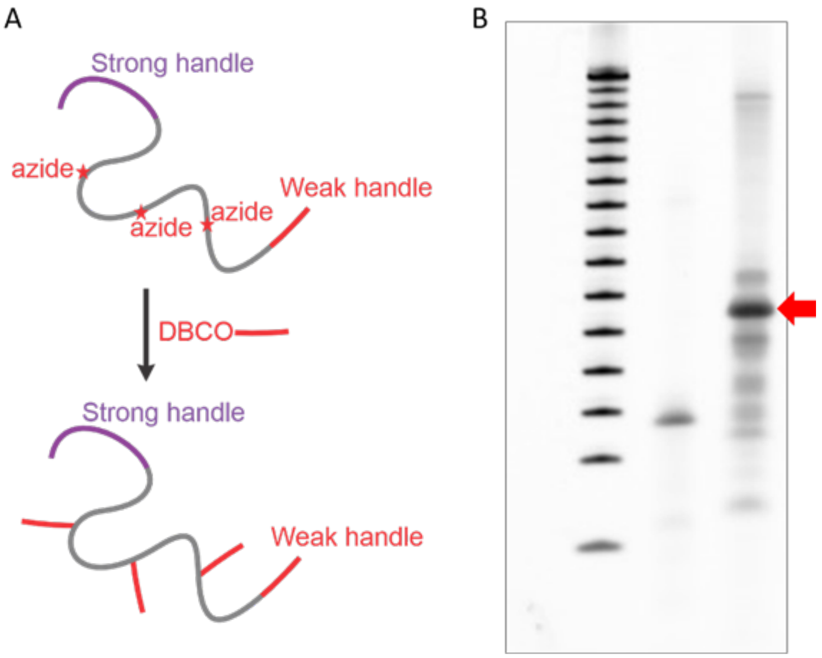


**Supplementary Figure 1.** Labeling of a nucleotide in each loop of G-quadruplex loop with a shearing handle. A) Schematic representation of the DBCO-azide click reaction to label shearing handles in each loop of the g-quadruplex. B) PAGE gel electrophoresis analysis to confirm the successful labeling reaction. Left lane: 50 bp ladder (Promega), Middle lane: ssDNA consisting of a G-quadruplex forming sequence, and right lane: ssDNA consisting of a G-quadruplex forming sequence with a shearing handle attached in each loop region of G-quadruplex. The red arrow indicates the expected product that was gel purified to use in DNC measurements.

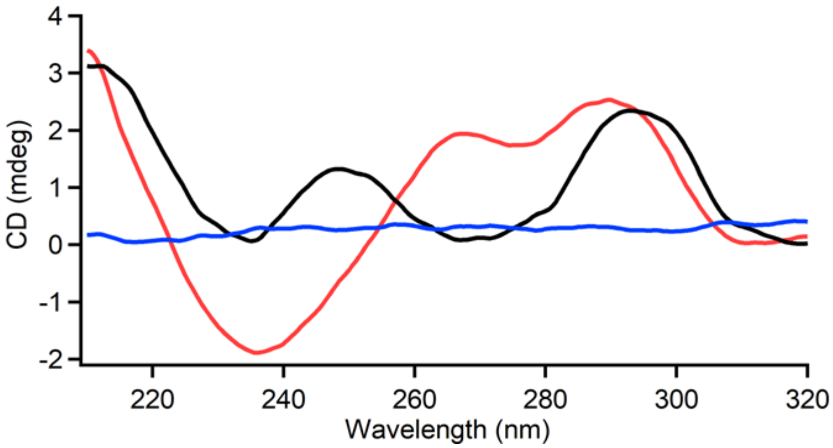


**Supplementary Figure 2.** Circular Dichroism spectroscopy of a human telomere G-quadruplex forming sequence in sodium ion (black), and potassium ion (red) containing buffer (10 mM Tris, pH 7.4).

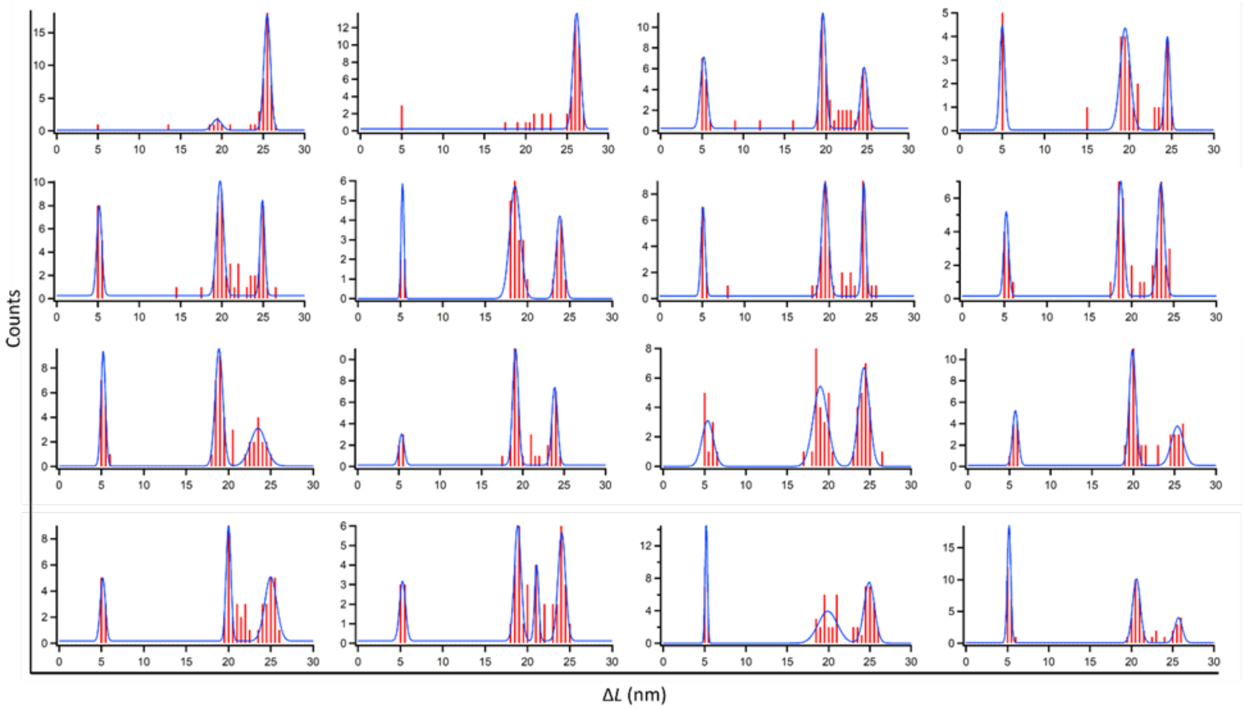


**Supplementary Figure 3.** Histograms show the distribution of the change-in-distances measured with g-quadruplex with the one weak shearing handle in a buffer consisting of sodium ions. Blue lines depict the multipeak Gaussian fitting. Three peaks indicate the identification of the three states of the target sequence; the Left peak: unfolding of the g-quadruplex, the middle peak: distance of the unfolded G-quadruplex, and right peak: end-to-end distance of the folded G-quadruplex.

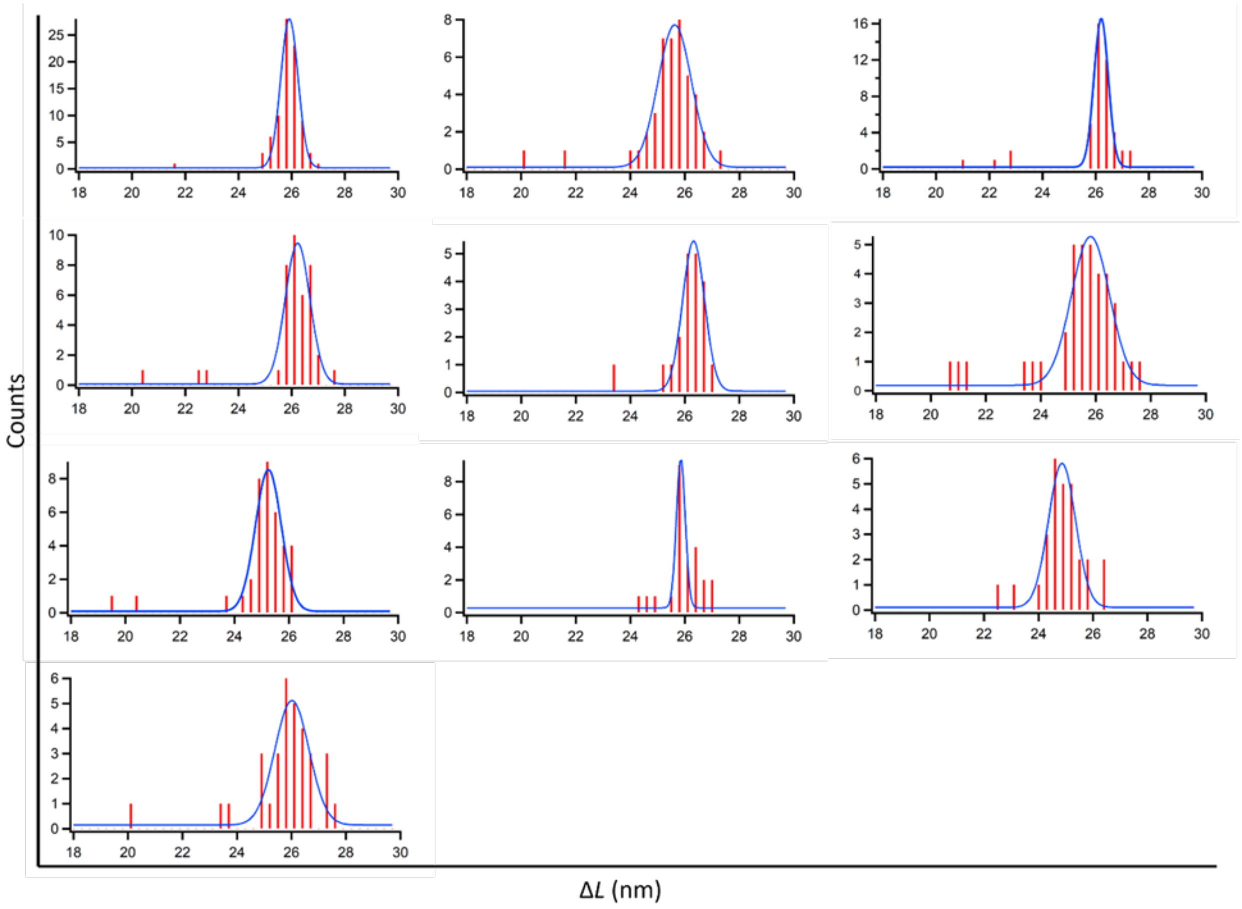


**Supplementary Figure 4.** Histograms show the distribution of the change-in-distances due to the unlooping of DNC consisting of a target G-quadruplex structure with one weak shearing handle in a buffer consisting of potassium ions. Blue lines depict the Gaussian fitting.

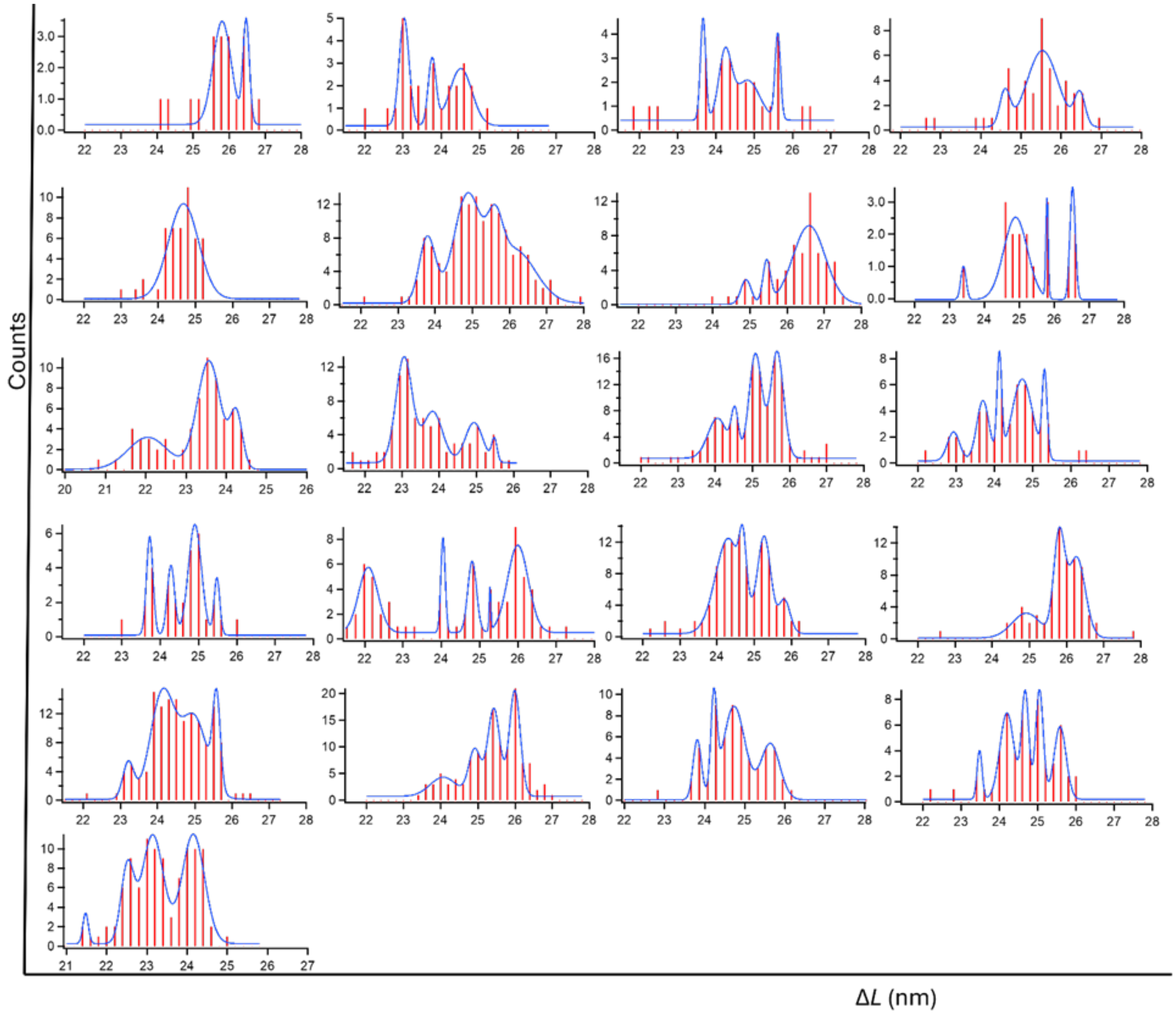


**Supplementary Figure 5.** Histograms show the distribution of the change-in-distance due to the unlooping of DNC to measure the distances in the individual G-quadruplexes formed in a buffer of potassium ions. Blue lines depict the multipeak Gaussian fitting.

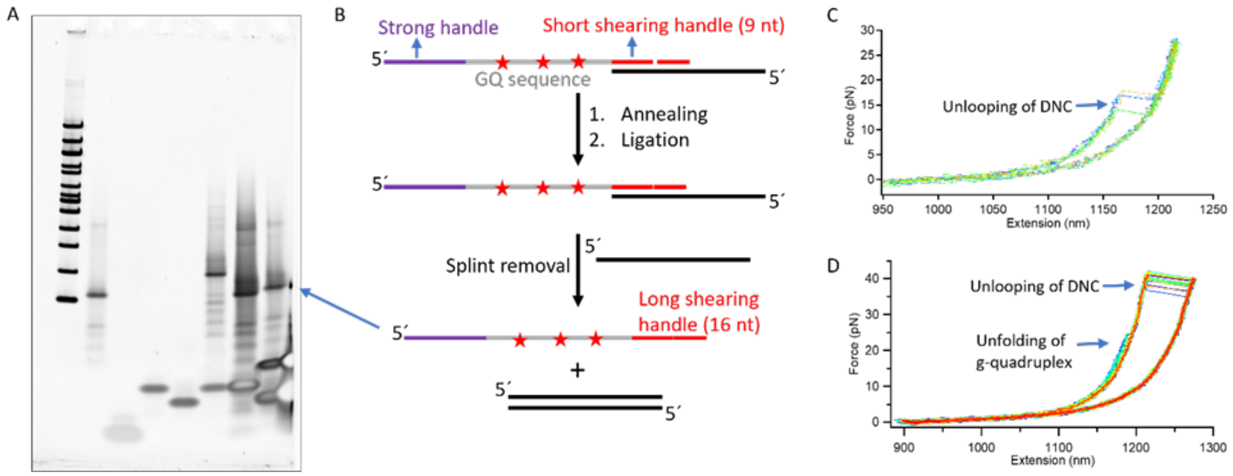


**Supplementary Figure 6.** Synthesis of g-quadruplex oligo with a longer shearing handle from the shorter shearing handle using DNA splint ligation. A) PAGE gel analysis (4-20% TBE gel, ThermoFisher Scientific) shows a successful synthesis of a target g-quadruplex with a long shearing handle (16 nt) from that of the short shearing handle (9 nt). B) Schematic representation of the strategy of the DNA splint ligation. The long blue arrow indicates the band of the desired product in (A) that was gel-purified to use in DNC measurements. Red stars indicate the position of incorporated azide in a loop nucleotide to label longer shearing handles by using DBCO-azide click chemistry as shown in supplementary figure 1. C) Force *vs* extension graph of DNC with target G-quadruplex with the short shearing handle that confirms the unlooping of DNC at a lower force than the unfolding of G-quadruplex. D) The Force vs extension graph of DNC with the target g-quadruplex with a long shearing handle confirms the unlooping of DNC at a higher force than the unfolding of the g-quadruplex.

**Supplementary Table 2.** 5´ to 3´ end-to-end distances measured in different types of DNA G-quadruplex conformations reported by NMR[1–5].

| **G-quadruplex conformations** | **Parallel** (PDB 1KF1) | **Antiparallel** (PDB 143D) | **Hybrid-1** (PDB 2HY9) | **Hybrid-2** (PDB 2JPZ) |
|---|---|---|---|---|
| 5´-3´ (nm) | 1.04 | 2.08 | 0.92 | 1.04 |

**Supplementary Table 3.** Distances between the 5´ end and the labeled residues measured in the different conformations of DNA g-quadruplex reported by NMR[1–5].

| Distances | Parallel (PDB 1KF1) | Antiparallel (PDB 143D) | Hybrid-1 (PDB 2HY9) | Hybrid-2 (PDB 2JPZ) |
|---|---|---|---|---|
| 5´-5´ loop (nm) | 1.89 | 1.87 | 1.07 | 1.79 |
| 5´- mid loop (nm) | 2.67 | 1.77 | 2.36 | 1.81 |
| 5´-3´ loop (nm) | 2.20 | 1.27 | 1.98 | 2.17 |
| 5´-3´ (nm) | 1.04 | 2.08 | 0.92 | 1.04 |

**Supplementary Table 4.** Theoretical estimation of the change-in-distance due to unfolding of g-quadruplex based on the anti-parallel conformation (PDB 143D)[1] in sodium ion as previously determined and unlooping of DNC.

| Unfolding directions | End-to-end distance measured in a folded conformation (NMR) (nm) | G-quadruplex $\Delta L_{unfold}$ (nm) | DNC $\Delta L_{unloop}$ (nm) |
|---|---|---|---|
| 5´ - 5´ loop | 1.87 | -0.07 | 68.99 |
| 5´ - mid loop | 1.77 | 2.28 | 65.81 |
| 5´- 3´ loop | 1.27 | 5.48 | 62.63 |
| 5´- 3´ | 2.08 | 7.38 | 59.89 |
